# The interaction between ORF18 and ORF30 is required for late gene expression in Kaposi’s sarcoma-associated herpesvirus

**DOI:** 10.1101/401976

**Authors:** Angelica F Castañeda, Britt A Glaunsinger

## Abstract

In the beta- and gammaherpesviruses, a specialized complex of viral transcriptional activators (vTAs) coordinate to direct expression of virus-encoded late genes, which are critical for viral assembly and whose transcription initiates only after the onset of viral DNA replication. The vTAs in Kaposi’s sarcoma-associated herpesvirus (KSHV) are ORF18, ORF24, ORF30, ORF31, ORF34, and ORF66. While the general organization of the vTA complex has been mapped, the individual roles of these proteins, and how they coordinate to activate late gene promoters, remains largely unknown. Here, we performed a comprehensive mutational analysis of the conserved residues in ORF18, which is a highly interconnected vTA component. Surprisingly, the mutants were largely selective for disrupting the interaction with ORF30 but not the other three ORF18 binding partners. Furthermore, disrupting the ORF18-ORF30 interaction weakened the vTA complex as a whole, and an ORF18 point mutant that failed to bind ORF30 was unable to complement an ORF18 null virus. Thus, contacts between individual vTAs are critical, as even small disruptions in this complex result in profound defects in KSHV late gene expression.

## Importance

Kaposi’s sarcoma-associated herpesvirus (KSHV) is the etiologic agent of Kaposi’s sarcoma and other B-cell cancers and remains a leading cause of death in immunocompromised individuals. A key step in the production of infectious virions is the transcription of viral late genes, which generates capsid and structural proteins and requires the coordination of six viral proteins that form a complex. The role of these proteins during transcription complex formation and the importance of protein-protein interactions are not well understood. Here, we focused on a central component of the complex, ORF18, and revealed that disruption of its interaction with even a single component of the complex (ORF30) prevents late gene expression and completion of the viral lifecycle. These findings underscore how individual interactions between the late gene transcription components are critical for both the stability and function of the complex.

## Introduction

A broadly conserved feature of the lifecycle of dsDNA viruses is that replication of the viral genome licenses transcription of a specific class of viral transcripts termed late genes. There is an intuitive logic behind this coupling, as late genes encode proteins that participate in progeny virion assembly and egress, and thus are not needed until newly synthesized genomes are ready for packaging. Additionally, late gene transcription requires ongoing DNA replication, and in the gammaherpesviruses Kaposi’s sarcoma-associated herpesvirus (KSHV) and Epstein-Barr virus (EBV) the increase in template abundance appears insufficient to explain the robust transcription of late genes whose products are required in large amounts (1).

While the mechanisms underlying late gene activation can vary across viral families, in the beta- and gammaherpesviruses, late gene promoters are strikingly minimalistic and primarily consist of a modified TATA box (TATT) and ~10-15 base pairs of variable flanking sequence (2–5). Despite this sequence simplicity, their transcription requires a dedicated set of at least six conserved viral transcriptional activators (vTAs) whose precise roles are only beginning to be uncovered. In KSHV, the vTAs are encoded by open reading frames (ORFs) 18, 24, 30, 31, 34, and 66 (6–15). The best characterized of the vTAs is a viral TATA-binding protein (TBP) mimic, encoded by ORF24 in KSHV, which binds both the late gene promoter and RNA polymerase II (Pol II) (4, 8, 16, 17). Beyond the viral TBP mimic, the only other vTA with a documented transcription-related function is pUL79 (homologous to KSHV ORF18) in human cytomegalovirus (HCMV), which promotes transcription elongation at late times of infection (18). Roles for the remaining vTAs remain largely elusive, although the KSHV ORF34 protein and its murine cytomegalovirus (MCMV) homolog pM95 may function as hub proteins, as they interact with numerous other vTAs (7, 9, 19). In addition to the six conserved vTAs, in EBV, the kinase activity of BGLF4 (homologous to KSHV ORF36) also contributes to the expression of late genes (20, 21).

Studies in both beta-and gammaherpesviruses indicate that the vTAs form a complex, the general organization of which has been mapped in MCMV and KSHV (7, 9, 10, 19) (Figure 1A). Notably, several recent reports demonstrate that specific interactions between the vTAs are critical for late gene transcription. In MCMV, mutation of conserved residues in pM91 (homologous to KSHV ORF30) that disrupt its interaction with pM79 (homologous to KSHV ORF18) renders the virus unable to transcribe late genes (19). Similarly, the interaction between KSHV ORFs 24 and 34 can be abrogated by a single amino acid mutation in ORF24 which prevents late gene transcription (9). Further delineating these contacts should provide foundational information relevant to understanding vTA complex function.

**Figure 1.**
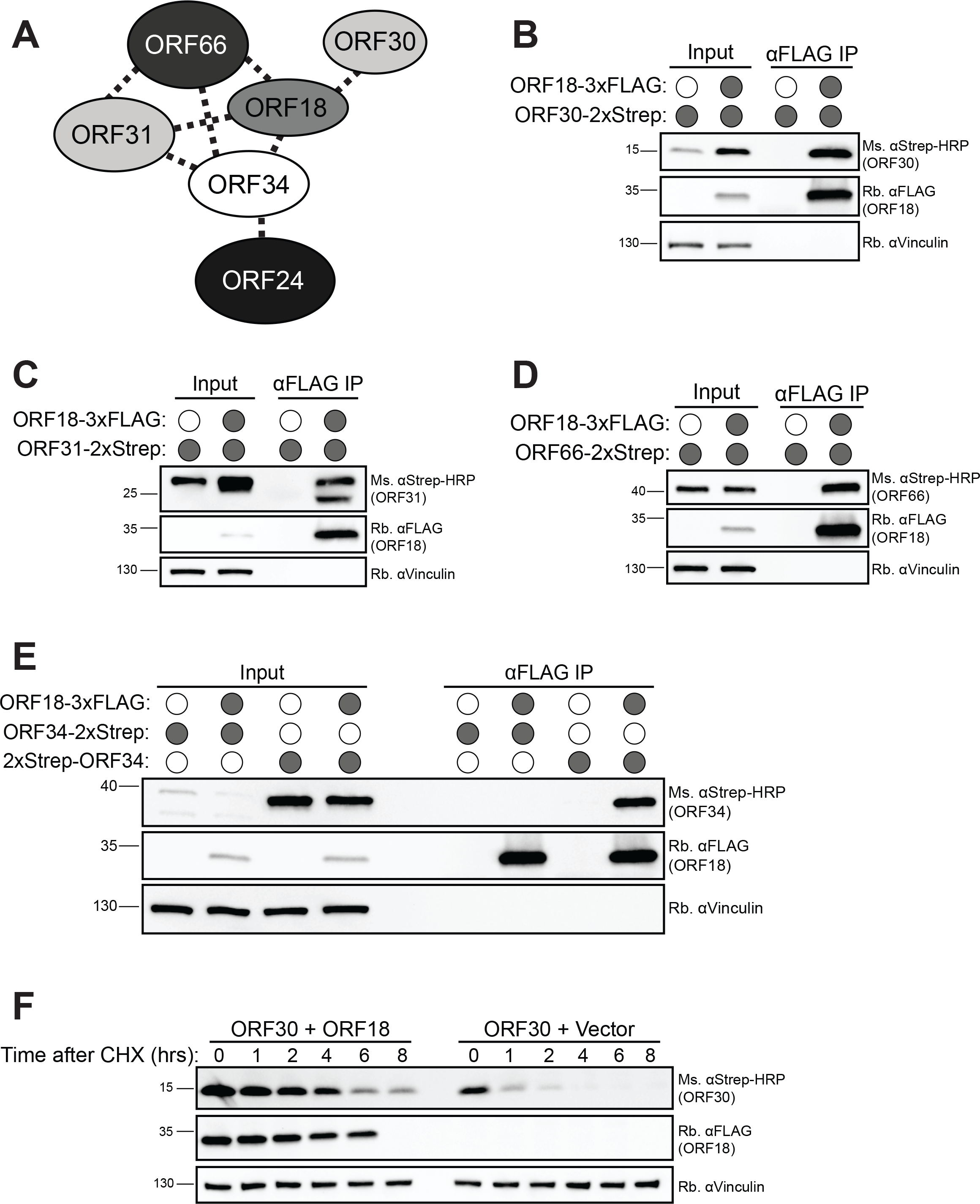
ORF18 interacts with ORFs 30, 31, 34, and 66. (A) Diagram of vTA interactions in KSHV from (9). (B-E) HEK293T cells were transfected with the indicated vTA plasmids, then subjected to co-IP using α-FLAG beads followed by western blot analysis with the indicated antibody to detect ORF18 and either ORF30 (*B*), ORF31 (*C*), ORF66 (*D*), and ORF34 (*E*). Input represents 2.5% of the lysate used for co-IP. Vinculin served as a loading control. (F) HEK293T cells were transfected with the indicated vTA plasmids. 24 h post transfection, cycloheximide was added to a final concentration of 100 μg/ml, and samples were collected at the indicated time points after the addition of cycloheximide. 25 μg of whole cell lysate was resolved using SDS-PAGE followed by western blot with the indicated antibodies. Vinculin served as a loading control.

The precise role of KSHV ORF18 in late gene transcription remains unknown, however it is predicted to interact with four of the five other vTAs (ORFs 30, 31, 34, and 66), suggesting that—like ORF34—it may play a central role in vTA complex organization (7, 9). Here, we performed an interaction screen of ORF18 mutants to comprehensively evaluate the roles of its conserved residues in mediating pairwise vTA binding. We reveal that ORF30 is particularly sensitive to mutation in ORF18, enabling isolation of mutants that selectively abrogate this interaction while retaining the contacts between ORF18 and the other vTAs. Disrupting the ORF18-ORF30 interaction not only prevents KSHV late gene transcription as measured by K8.1 expression, but also appears to weaken assembly of the remaining vTA complex. These findings underscore the key role that ORF18 plays in late gene transcription and suggest that disrupting just one of its interactions has a destabilizing effect on the vTA complex as a whole.

## Results

### ORF18 interacts with ORF30, ORF31, ORF34, and ORF66

Previous work using a split luciferase-based interaction screen suggested that ORF18 is highly interconnected with other viral late gene activators, as it interacted with the majority of the proteins in the viral transcription pre-initiation complex (vPIC) (9). To independently confirm the binding of ORF18 to ORFs 30, 31, 34, and 66 in the absence of other viral factors, we assessed its ability to co-immunoprecipitate (co-IP) with each of these vTAs in transfected HEK293T cells. Consistent with the screening data, ORF18-3xFLAG interacted robustly with C-terminal 2xStrep-tagged versions of ORF30, ORF31, and ORF66 (Figure 1B, C, D). Although ORF18-3xFLAG did not interact with ORF34 tagged on its C-terminus, the interaction was recovered upon moving the 2xStrep tag to the N-terminus of ORF34 (Figure 1E). Furthermore, we consistently observed that the expression of ORF30 was higher when co-expressed with ORF18 than when transfected with vector control, suggesting that ORF18 may stabilize ORF30 (Figure 1B). To determine whether ORF18 had a stabilizing effect on ORF30, the half-life of ORF30 was measured in both the presence and absence of ORF18. As can be seen in Figure 1F, ORF30 stability increased significantly upon co-expression with ORF18.

### An interaction screen of ORF18 mutants reveals the role of conserved residues in interactions with the other vTAs

To evaluate the importance of the interaction between ORF18 and its individual vTA contacts, we aimed to identify point mutations that disrupted binding to individual vTAs but did not destroy the integrity of the complex. Since the late gene vTA complex is conserved across the beta- and gammaherpesviruses we reasoned that the individual points of contact might depend on conserved amino acid residues. We performed a multiple sequence alignment between KSHV ORF18 and its homologs in five other beta- and gamma herpesviruses (MHV68 ORF18, HCMV pUL79, MCMV pM79, EBV BVLF1, and BHV ORF18) using MUSCLE (22). The equence alignment revealed 25 single conserved residues, including six pairs of adjacent conserved residues, which are depicted in Figure 2A as a schematic of the primary structure of ORF18 showing the positions of conserved residues. We mutated each of the 25 conserved residues to alanine in ORF18-3xFLAG and made double alanine mutations in the six cases of adjacent conserved residues (Figure 2A). Each of these 31 mutants was screened individually for the ability to interact with ORFs 30, 31, and 66 by co-IP followed by western blot (data not shown). To account for differences in expression between the ORF18 mutants, we calculated the co-IP efficiency of each of the mutants, as described in the methods. These data were used to generate a heat map, which displays the pair-wise interaction efficiencies on a color scale where lighter blocks represent reduced binding and darker blocks represents increased binding relative to WT (Figure 2B). Overall, the data revealed that ORF30 was the most sensitive to mutations in ORF18, as 24 out of the 31 mutants displayed reduced or no binding. ORF31 showed variable sensitivity to ORF18 mutation (with some mutants even increasing the interaction efficiency), while the ORF66-ORF18 interaction was relatively refractory to the ORF18 point mutations (Figure 2B).

**Figure 2.**
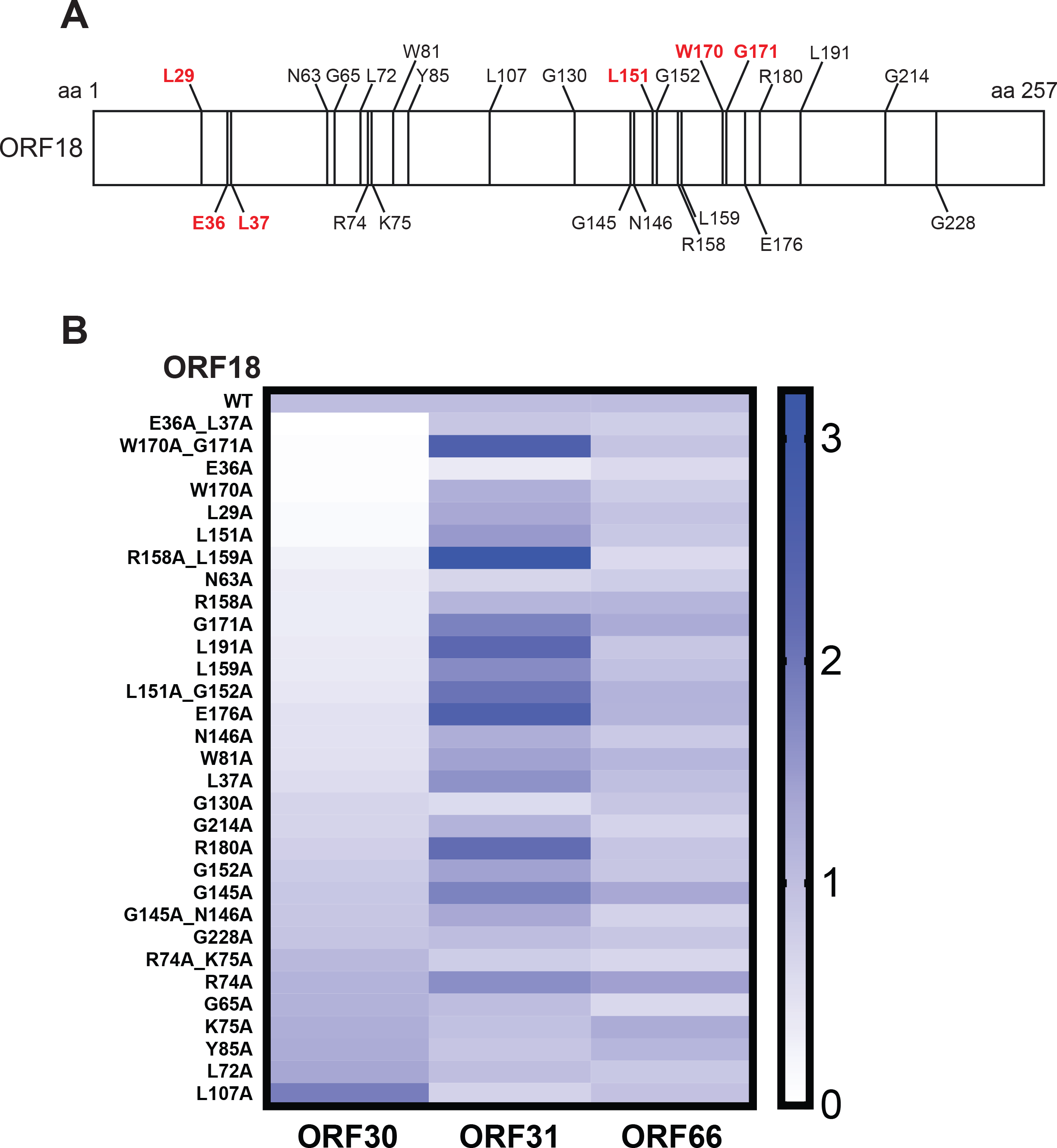
ORF18 mutant screen for interactions with ORFs 30, 31, and 66. (A) Diagram depicting the conserved residues in KSHV ORF18 in a MUSCLE alignment with MHV68 ORF18, EBV BVLF1, HCMV pUL79, MCMV pUL79, and BHV4 ORF18. Red color denotes amino acids found to have interactions <10% of WT with any of the vTAs. (B) Heat map of the co-IP efficiency of each ORF18 mutant against ORFs 30, 31, and 66.

We focused on the six ORF18 mutants that exhibited <10% co-IP efficiency, relative to WT, with any vTA in our initial screen (L29A, E36A, L151A, W170A, E36A_L37A, and W170A_G171A; highlighted in red in Figure 2A). These were re-screened 3-4 independent times in co-IP assays with vTA components ORFs 30, 31, 34, and 66, and the co-IP efficiencies were calculated as described in the methods then plotted relative to values obtained for WT ORF18 (Figure 3A-D). All six of these ORF18 mutants had severe defects in their ability to co-IP ORF30, but none were consistently different than WT ORF18 for interaction with ORFs 31, 34, and 66 (Figure 3B-D). Among the six mutants, ORF18^E36A_L37A^ and ORF18^W170A_G171A^ showed no detectable binding to ORF30, with ORF18^E36A_L37A^ retaining near-WT levels of interaction with the remaining vTAs.

**Figure 3.**
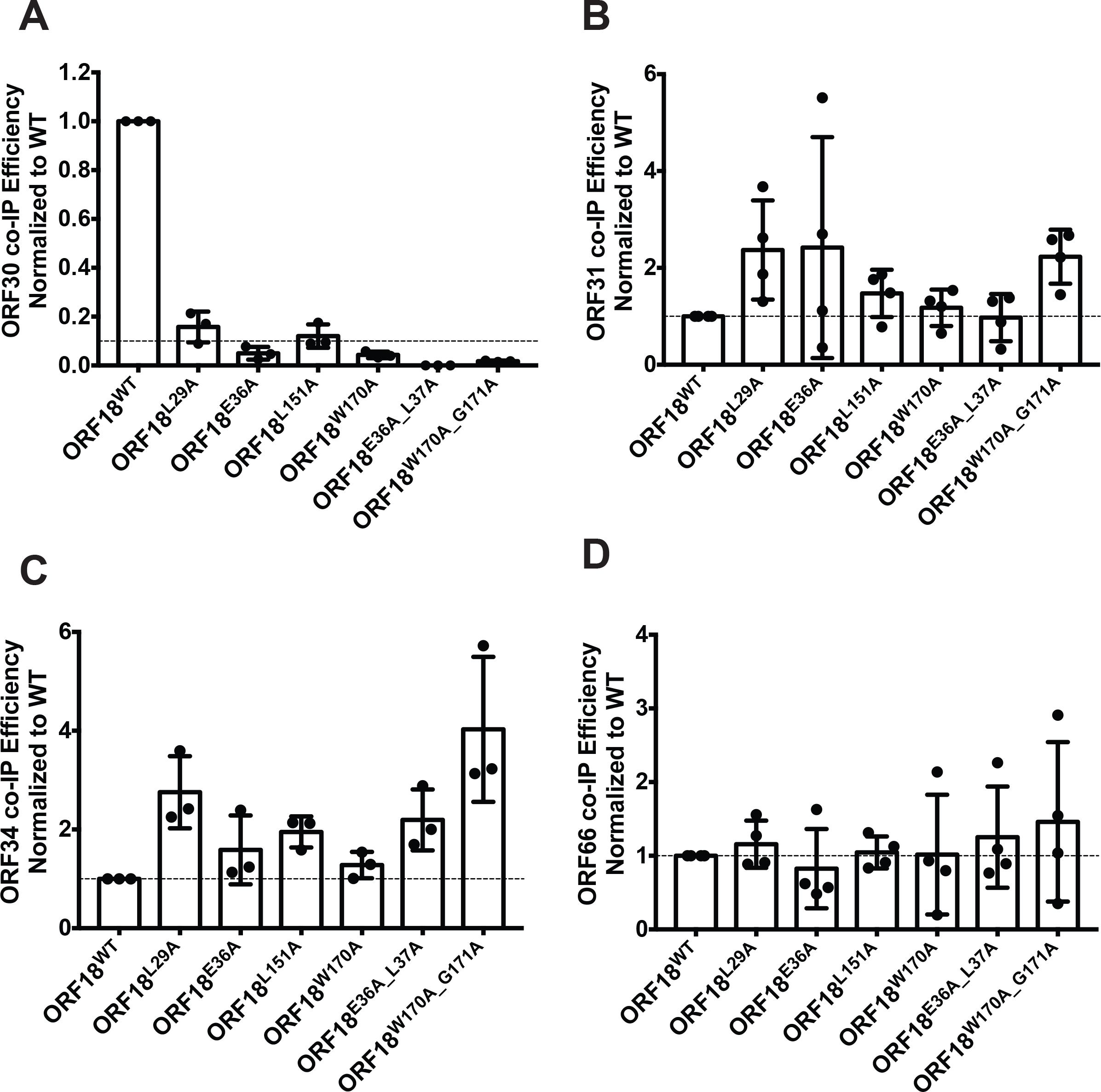
Six ORF18 mutants are consistently defective for interaction with ORF30. (A-D) HEK293T cells were transfected with the indicated vTA plasmids, then subjected to co-IP using α-FLAG beads followed by western blot analysis to detect the ability of WT or mutant ORF18 to interact with ORF30 (*A*), ORF31 (*B*), ORF34 (*C*), and ORF66 (*D*). The co-IP efficiency was calculated as described in the text for 3-4 independent experimental replicates and plotted as bar graphs. In *(A)*, the dotted line represents Y = 0.1, and in *(B-D)*, the dotted line represents Y = 1.0.

### ORF18 point mutants that weaken the interaction between ORF18 and ORF30 have a reduced capacity to activate the K8.1 late gene promoter

A reporter assay has been developed in which the co-expression of the six individual vTAs can specifically activate a KSHV (or EBV) late gene promoter in transfected HEK293T cells (9, 10). We used this assay as an initial proxy for the ability of the six ORF18 mutants described above to activate late gene transcription. HEK293T cells were co-transfected with each of the vTAs, including either WT or mutant ORF18, and firefly luciferase reporter plasmids driven by either the late K8.1 promoter or, as a control, the early ORF57 promoter (Figure 4A). A plasmid containing constitutively expressed Renilla luciferase was co-transfected with each sample to normalize for transfection efficiency. As expected, inclusion of WT ORF18 with the remaining vTA complex resulted in specific activation of the K8.1 late promoter, but not of the early ORF57 promoter (Figure 4B). ORF18 mutants L29A, E36A, and L151A modestly reduced activation of the late promoter, whereas more significant defects were observed with mutants W170A, E36A_L37A, and W170A_G171A (Figure 4B). Although ORF18^W170A_G171A^ had the most pronounced transcriptional defect, this mutant showed somewhat more variability than ORF18^E36A_L37A^ in its interactions with the other vTA components (Figures 3 & 4). Thus, we considered ORF18^E36A_L37A^ to be the top candidate for selectively analyzing the importance of the ORF18-ORF30 interaction for KSHV late gene transcription.

**Figure 4.**
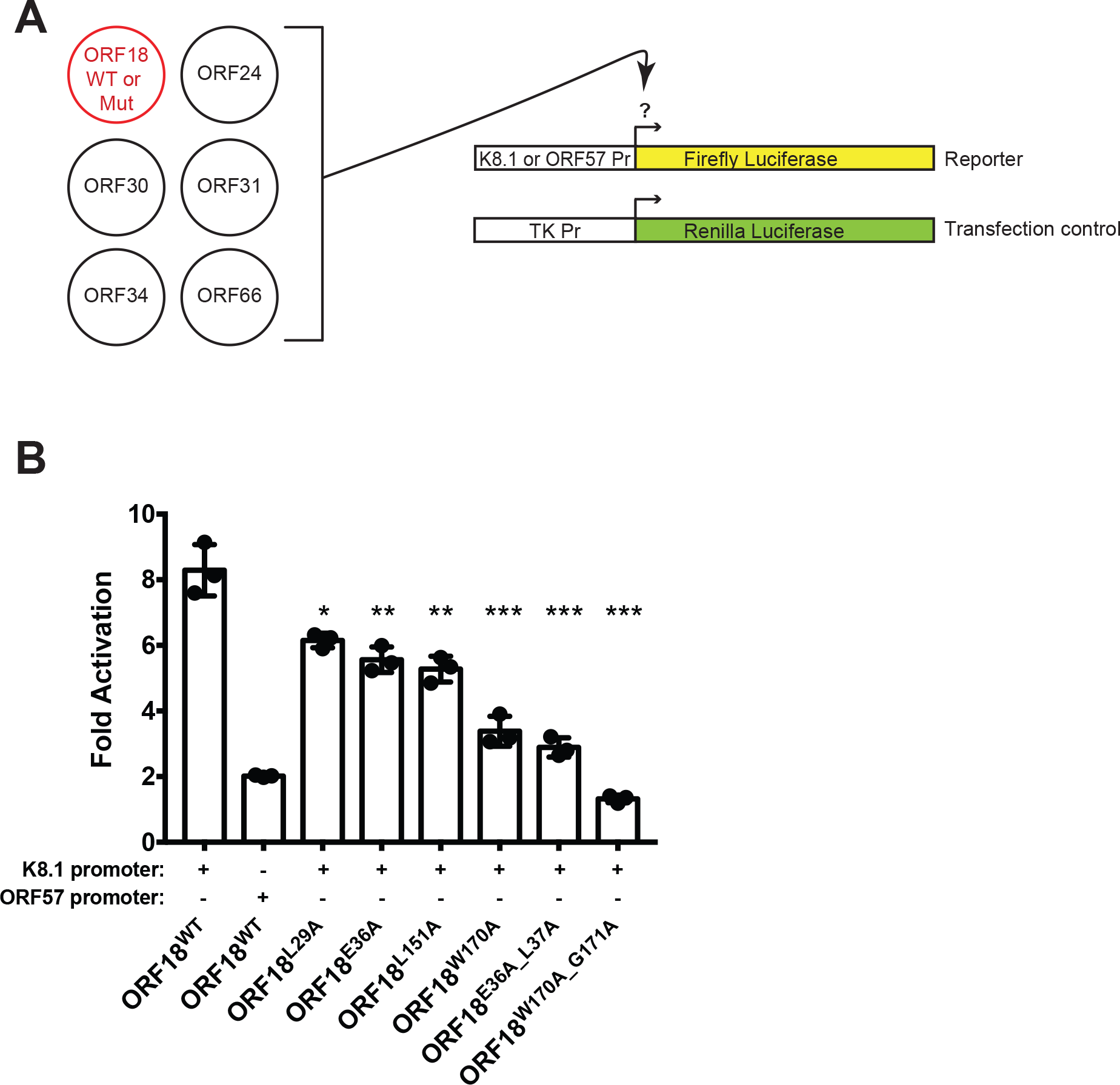
Impairing the interaction between ORF18 and ORF30 reduces activation of the late K8.1 promoter. (A) Diagram depicting the vector combinations that were transfected for the late gene reporter assay. (B) HEK293T cells were transfected with the vTA plasmids including WT or mutant ORF18, the K8.1 or ORF57 promoter reporter plasmids, and the pRL-TK Renilla plasmid as a transfection control. 24 h post-transfection the lysates were harvested and luciferase activity was measured. Data shown are from 3 independent biological replicates, with statistics calculated using an unpaired t-test where (*) p < 0.05, (**) p < 0.005, and (***) p < 0.0007.

### The interaction between ORF18 and ORF30 affects assembly of the vTA complex

The transcriptional defect of the ORF18^E36A_L37A^ mutant in the reporter assay could be due to a defect in assembly of the complex or due to defects in downstream events. To distinguish between these possibilities, WT or mutant ORF18-3xFLAG was co-transfected into HEK293T cells with each of the other Strep-tagged vTAs. We then performed an α-FLAG IP, revealing that purification of WT ORF18 led to co-IP of the complete vTA complex including Pol II, which has been shown to interact with ORF24 in KSHV (8) (Figure 5). We noted that in this assay ORF18^E36A_L37A^ was more weakly expressed than WT ORF18, so to compare complex formation with equivalent amounts of each protein, we titrated down the amount of WT ORF18 to match the levels of ORF18^E36A_L37A^. Similar to our observation in a pairwise co-IP (Figure 1B), the ORF30 protein abundance decreased as the expression of ORF18 was reduced (Figure 5); however, the complete vTA complex still co-purified even with reduced levels of WT ORF18 (Figure 5). Notably, the vTA complex was recovered at lower levels in the presence of ORF18^E36A_L37A^. When imaged at a longer exposure, all of the vTAs, with the exception of ORF30, remained associated with ORF18^E36A_L37A^ (Figure 5, far right panel). Thus, the selective loss of the ORF18-ORF30 interaction may reduce the overall stability of the vTA complex.

**Figure 5.**
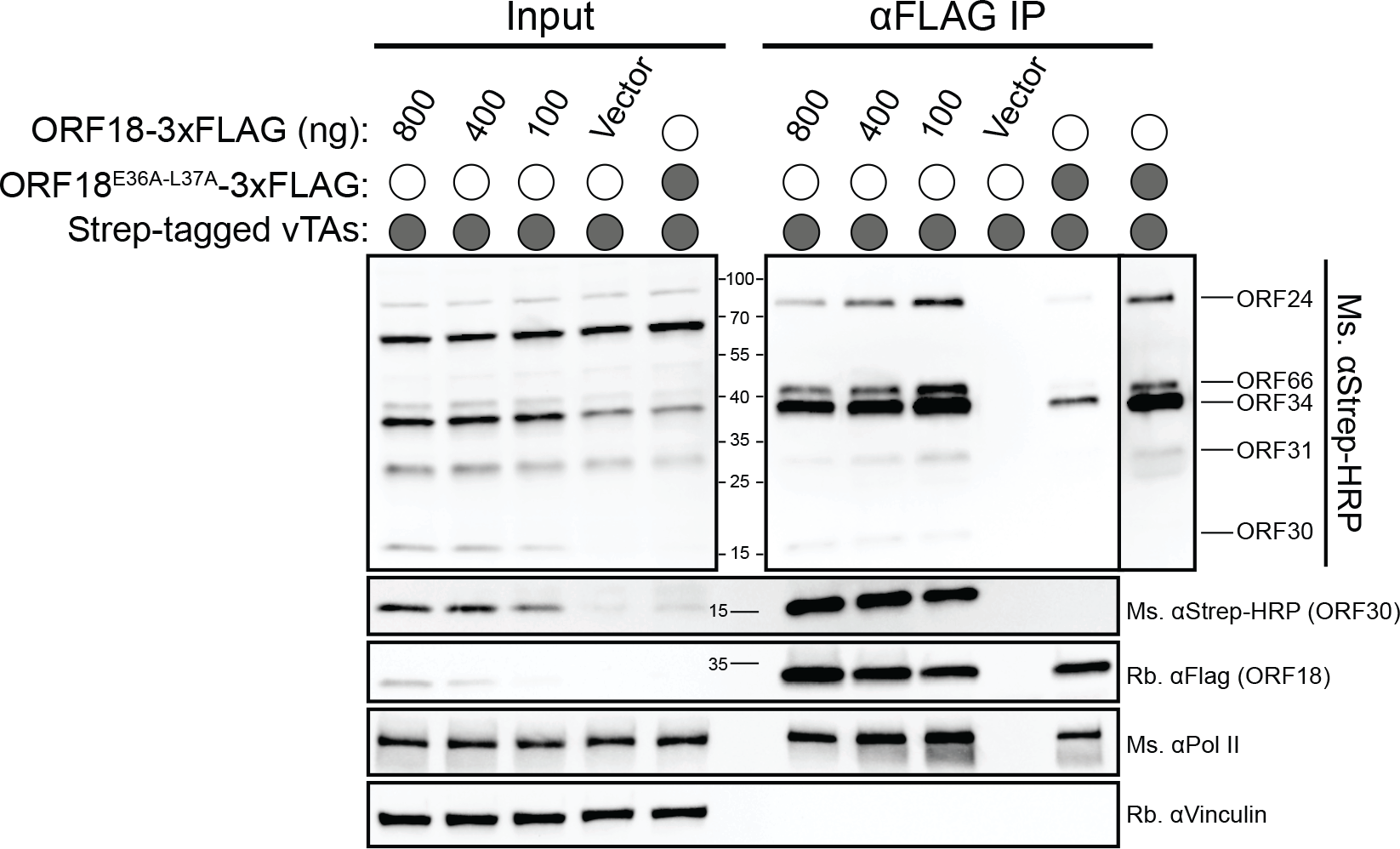
Disrupting the interaction between ORF18 and ORF30 weakens the assembly of the vTA complex. HEK293T cells were transfected with the indicated vTA plasmids then subjected to co- IP using α-FLAG beads followed by western blot analysis with the indicated antibody. Far right boxed lane is a longer exposure of the ORF18^E36A_L37A^ IP lane. Input represents 2.5% of the lysate used for co-IP. Vinculin was used as a loading control.

### The interaction between ORF18 and ORF30 is crucial for expression of the late gene K8.1

Next, to characterize the effect of ORF18^E36A_L37A^ on the viral replication cycle, we tested the ability of this mutant to complement the late gene expression defect of a KSHV mutant lacking ORF18 (18.stop) (6). The renal carcinoma cell line iSLK harbors the virus (either WT or 18.stop) in a latent state, which can be reactivated upon expression of the doxycycline-inducible major lytic transactivator RTA and treatment with sodium butyrate. Using lentiviral transduction, we generated stable, doxycycline-inducible versions of the 18.stop iSLK cells expressing either ORF18-3xFLAG or ORF18^E36A_L37A^-3xFLAG (18.stop.ORF18^WT^ and 18.stop.ORF18^E36A_L37A^, respectively). The cells were assayed 72 hours post lytic reactivation for their ability to replicate DNA, express early and late proteins, and produce progeny virions. Although we observed a modest decrease of viral DNA replication in the 18.stop cells, as measured by qPCR, upon complementation with either WT ORF18 or ORF18^E36A_L37A^, the levels of DNA replication were not significantly different from iSLK cells infected with WT KSHV (Figure 6A). Notably, the 18.stop.ORF18^E36A_L37A^ cell line expressed more ORF18 than the 18.stop.ORF18^WT^ cell line, in contrast to the reduced expression of the mutant in HEK293T cells (Figure 6B, compare to levels in Figure 5). However, while both reactivated 18.stop.ORF18^WT^ and 18.stop.ORF18^E36A_L37A^ cell lines expressed equivalent levels of the ORF59 early protein, only 18.stop.ORF18^WT^ was able to rescue expression of the model late gene K8.1 (Figure 6B).

**Figure 6.**
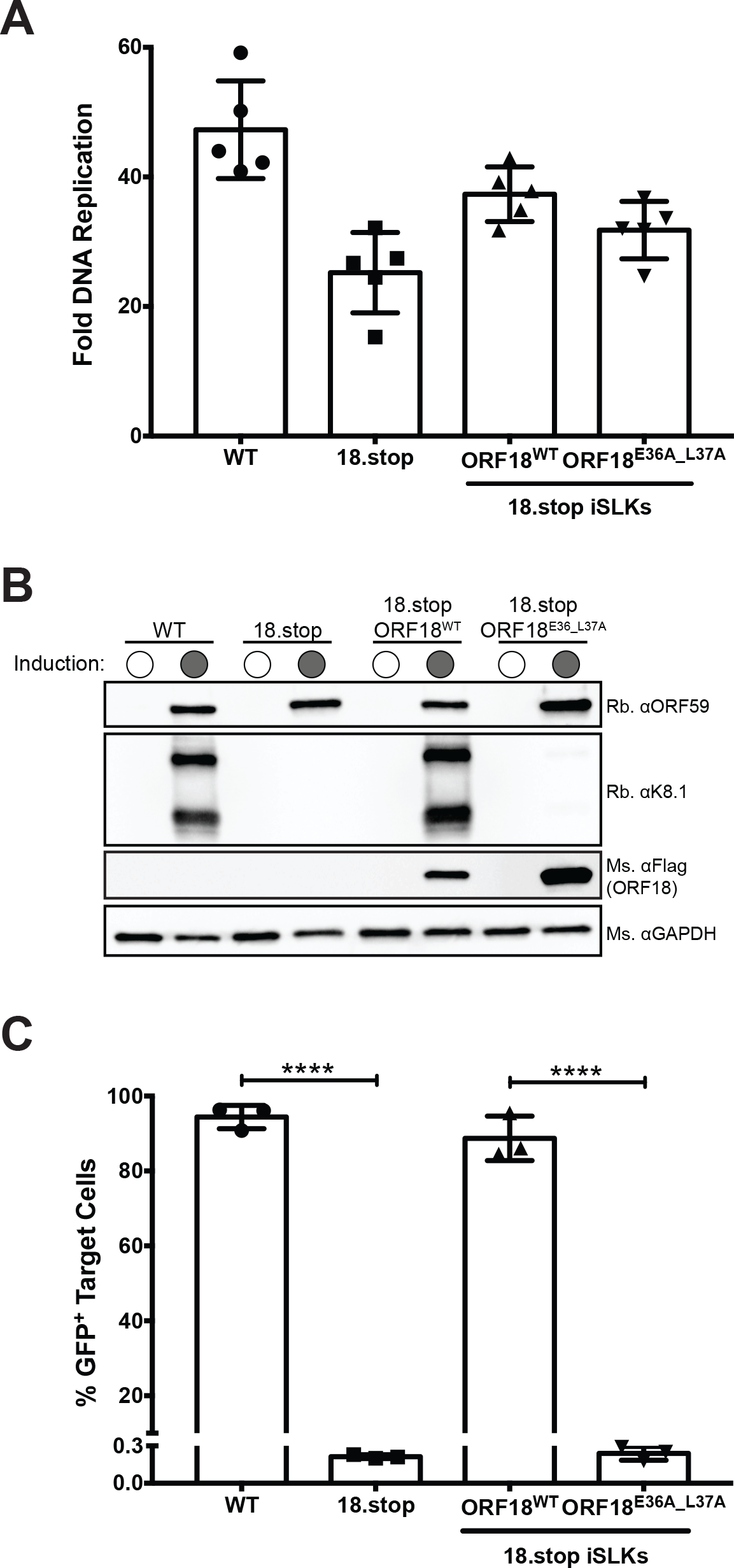
Characterizing the effect of the E36A_L37A ORF18 mutation on the virus. (A) iSLK cells latently infected with WT KSHV, 18.Stop KSHV, or 18.Stop complemented with ORF18^WT^ or ORF18^E36A_L37A^ were induced to enter the lytic cycle with 1 μg/ml doxycycline and 1 mM sodium butyrate. 72 h post induction, DNA was isolated and fold viral DNA replication was measured by qPCR before and after induction of the lytic cycle. Data shown are from 5 independent biological replicates. (B) Western blots of the expression of the early protein ORF59, the late protein K8.1, ORF18^WT^, and ORF18^E36A_L37A^ in the indicated cell lines induced as described in (*A*). GAPDH was used as a loading control. (C) HEK293T target cells were spinfected with filtered supernatant from induced cells. Progeny virion production was measured 24 h after supernatant transfer by flow cytometry of GFP+ target cells. Data shown are from 3independent biological replicates with statistics calculated using an unpaired t-test where (****) p < 0.0001.

We then evaluated the level of KSHV virion production from the parental WT KSHV-infected iSLK cells, as well as from the 18.stop, 18.stop.ORF18^WT^, and 18.stop.ORF18^E36A_L37A^ cell lines using a supernatant transfer assay. KSHV produced from iSLK cells contains a constitutively expressed GFP, enabling quantitation of infected recipient cells by flow cytometry (23). Consistent with its late gene expression defect, neither the 18.stop nor the 18.stop.ORF18^E36A_L37A^ cell lines were able to produce progeny virions, whereas virion production in the 18.stop.ORF18^WT^ cells was indistinguishable from the WT KSHV-infected iSLK cells (Figure 6C). Collectively, these data demonstrate that the specific interaction between ORF18 and ORF30 is essential for K8.1 late gene expression and virion production during KSHV infection.

### The analogous mutation in HCMV pUL79 disrupts its interaction with pUL91

As shown in Figure 7A, the E36_L37 residues are conserved across the beta- and gammaherpesvirus ORF18 homologs. To determine whether these amino acids are similarly important in a betaherpesvirus, we engineered the corresponding double mutation in HCMV pUL79 (pUL79^E48A_L49A^-3xFLAG). Similar to our observation with KSHV ORF30, HCMV pUL91 protein expression was significantly decreased in the absence of its WT pUL79 binding partner (Figure 7B). This is consistent with the idea that pUL79 binding stabilizes pUL91. Furthermore, in co-IP assays we detected a robust interaction between pUL79 and pUL91, which was impaired in the presence of pUL79^E48A_L49A^, even when we accounted for the expression level differences of pUL91 (Figure 7B). This suggests that the overall protein-protein interface may be conserved in the analogous ORF18-ORF30 interaction across the beta- and gammaherpesviruses.

**Figure 7.**
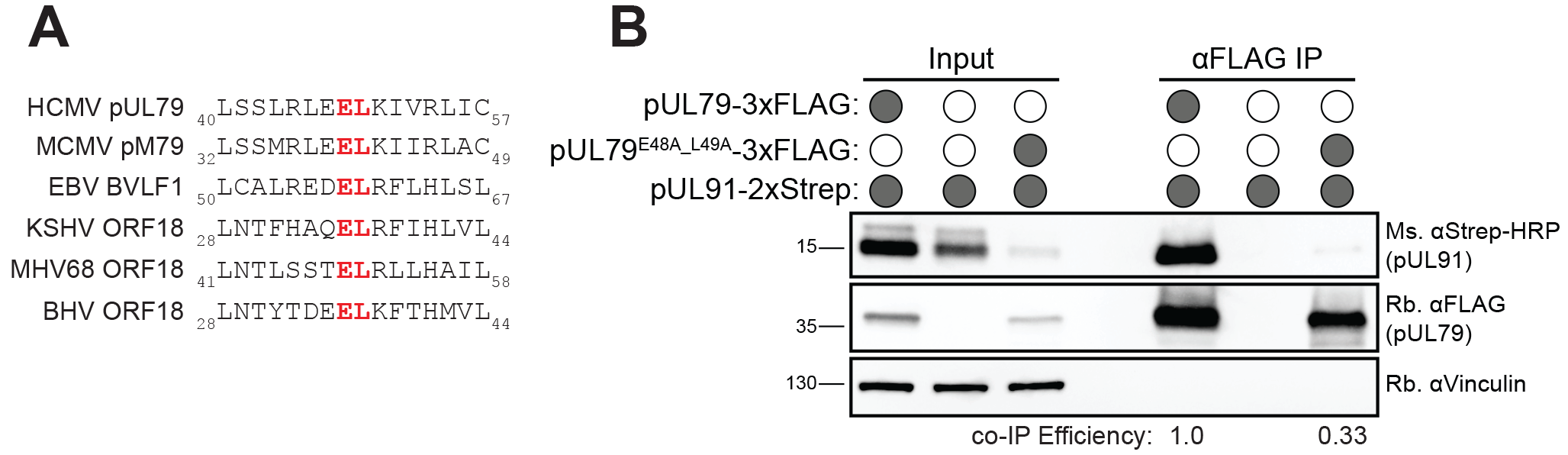
Mutating E48_L49 in pUL79 disrupts its interaction with pUL91. (A) MUSCLE multiple sequence alignment for HCMV pUL79 and homologs showing the location of conserved amino acids that correspond to E36_L37 in KSHV ORF18. (B) HEK293T cells were transfected with the indicated plasmids, then subjected to co-IP using α-FLAG beads followed by western blot analysis with the indicated antibody. To normalize for the difference in expression of pUL91 in the different transfection conditions, the co-IP efficiency was multiplied by the fold expression of pUL91 in the presence of WT pUL79 versus E48A_L49A pUL79. Input represents 2.5% of lysate used for co-IP. Vinculin was used as a loading control.

## Discussion

Elucidating the architecture of the six-member vPIC complex is central to understanding the mechanism underlying viral late gene expression in beta- and gammaherpesviruses. Although their functions are largely unknown, each of these viral transcription regulators is essential for late gene promoter activation and evidence increasingly suggests that their ability to form a complex is crucial for transcriptional activity (9, 10, 19). Here, we reveal that selective disruption of an individual protein-protein contact between KSHV ORF18 and ORF30 within the vPIC is sufficient to abrogate K8.1 late gene expression and virion production in infected cells, emphasizing the sensitivity of the complex to organizational perturbation.

We selected ORF18 for mutational screening due to its ability to form pairwise interactions with the majority of other vPIC components, which suggested that it might serve as an organizational hub for vPIC assembly, similar to what has been proposed for ORF34 (7, 9). However, the 31 tested mutants of ORF18 revealed that conserved residues across the length of the protein are extensively—and largely selectively— required for ORF30 binding. This is in contrast to the interaction between the vPIC components ORF24 and ORF34, where the interaction can be localized to a 17 amino acid stretch of ORF24 (9). The observation that the majority of the ORF18 point mutations that disrupt the interaction with ORF30 do not affect its binding to ORFs 31, 34, and 66 indicates that these mutations do not significantly alter the overall folding or structure of ORF18. In MCMV, the organization of the vTA complex is similar to KSHV, except that pM92 (homologous to KSHV ORF31) interacts with pM87 (homologous to KSHV ORF24), whereas in KSHV this interaction is bridged through ORF34 (19). Our data complement recent findings in MCMV, in which mutations of the ORF30 homolog (pM91) that perturb the interaction with the ORF18 homolog (pM79) similarly cause a defect in the expression of late genes (19). Thus, mutations that disrupt the ORF18-ORF30 protein-protein interface in either protein cause the same phenotype.

The ORF18-ORF30 interaction appears sensitive to single amino acid changes in the protein-protein interface. The ORF18^E36A_L37A^ mutation does not impair binding to its other vPIC partners in pairwise co-IP experiments and ORF30 does not engage in pairwise interactions with vPIC components other than ORF18 (7, 9). It is therefore notable that the efficiency of the vPIC complex assembly is reduced in HEK293T cells in the presence of the ORF18^E36A_L37A^ mutant, suggesting that the ORF18-ORF30 interaction contributes to the stability of the complex as a whole. The interaction between ORF18 and ORF30 may change the conformation of ORF18, allowing it to interact more strongly with the other vTAs. Alternatively, the E36A_L37A mutation may contribute to binding defects between ORF18 and the other vTAs that are not observed with pairwise interactions, but are enhanced in the presence of all the ORF18 binding partners.

We observed that ORF30 protein expression was consistently higher in the presence of WT ORF18 or ORF18 mutants that retained ORF30 binding, suggesting that ORF18 helps stabilize ORF30. *In silico* protein stability prediction studies have suggested that protein stability is in part affected by protein length, where proteins that are less than 100 amino acids tend to be less stable (24, 25). One explanation for the higher expression of ORF30 in the presence of ORF18 could therefore be that the 77 amino acid ORF30 is protected from degradation by ORF18. Another possibility is that ORF18 helps keep ORF30 correctly folded—this has been proposed as a mechanism that stabilizes proteins which have interaction partners (26). The interaction-induced stability of a protein often correlates with the relative concentration of its binding partners (26), as we observed when we titrated down the amount of ORF18 in the context of the complete vTA complex. A similar observation has been made between KSHV proteins ORF36 and ORF45, where ORF36 was dependent on the interaction with ORF45 for stabilization (27). We did not observe a similar correlation with levels of the other ORF18- associated vPIC proteins, and thus their stability may not require protective interactions. We also observed a stabilizing effect of pUL79 on pUL91, indicating this may be a conserved feature of this interaction in other beta- and gammaherpesviruses.

The fact that the late gene expression defect of the ORF18^E36A_L37A^ mutant is exacerbated in the context of the virus, compared to in the plasmid promoter activation assay, likely reflects the fact that the plasmid assay measures basal promoter activation but misses other regulatory components of this cascade. For example, the origin of lytic replication is required *in cis* for late gene expression in related gammaherpesviruses (3, 28) and the reporter assay does not capture the important contribution of viral DNA replication towards late gene expression. This may explain why some mutants that are defective for ORF30 binding (e.g. L29A, E36A, L151A) retain partial plasmid promoter activity; perhaps some weak binding between ORF18 and ORF30 enables basal activation of the promoter in a context where the vPIC components are overexpressed. Alternatively, some of the mutations may cause ORF18 to bind to ORF30 more transiently, but their vPIC interaction becomes stabilized in the presence of a late gene promoter.

In summary, the absence of K8.1 late gene expression in the KSHV ORF18.stop-infected cells complemented with ORF18^E36A_L37A^ may derive from a cascade of phenotypes: the failure to recruit ORF30 to the vPIC, the ensuing reduction in the efficiency of overall vPIC complex assembly, and the reduced stability of ORF30 (if it also has additional vPIC-independent functions). Ultimately, generating antibodies that recognize the endogenous KSHV vPIC components will enable these phenotypes to be explored further during infection. In addition, information about the 3-dimensional structure of the vPIC would significantly enhance our understanding of this unique transcription complex, as it is becoming increasingly clear that even small disruptions to the complex dramatically impact completion of the viral lifecycle.

## Materials and Methods

### Plasmids and Plasmid construction

To generate ORF18-3xFLAG pCDNA4, ORF18 was subcloned into the BamHI and NotI sites of pCDNA-3xFLAG. The point mutations in ORF18 were generated using two primer site-directed mutagenesis with Kapa HiFi polymerase (Roche) with primers 1-62 listed in Table 1. All subsequent plasmids described below were generated using InFusion cloning (Clontech) unless indicated otherwise. To generate plasmid pLVX-TetOneZeo, zeocin resistance was PCR amplified out of plasmid pLJM1-EGFP-Zeo with primers 63/64 (Table 1) and used to replace the puromycin resistance in pLVX-TetOne™-Puro (Clontech) using the AvrII and MluI restriction sites. To generate pLVX-TetOneZeo-ORF18^WT^-3xFLAG and pLVX-TetOneZeo-ORF18^E36A_L37A^-z3xFLAG, ORF18^WT^-3xFLAG and ORF18^E36_37A^-3xFLAG were PCR amplified from each respective pCDNA4 plasmid using primers 71/72 and inserted into the EcoRI and BamHI sites of pLVX-TetOne-Zeo. To generate UL79-3xFLAG pCDNA4 and UL91-2xStrep pCDNA4, UL79 was PCR amplified with primers 67/68 (Table 1) and UL91 was PCR amplified with primers 69/70 (Table 1) from HCMV Towne strain, which was kindly provided by Dr. Laurent Coscoy, and cloned into the BamHI and NotI sites of 3xFLAG (Cterm) pCDNA4 or 2xStrep (Cterm) pCDNA4. UL79^E48A_L49A^ was generated with two primer site-directed mutagenesis using Kapa HiFi polymerase with primers 73/74 (Table 1). To make 2xStrep-ORF34 pCDNA4, ORF34 was PCR amplified from ORF34-2xStrep pCDNA4 with primers 65/66 and cloned into the NotI and XbaI sites of 2xStrep (Nterm) pCDNA4. Plasmid K8.1 Pr pGL4.16 contains the minimal K8.1 promoter and ORF57 Pr pGL4.16 contains a minimal ORF57 early gene promoter and have been described previously (9). Plasmids ORF18-2xStrep pCDNA4, ORF24-2xStrep pCDNA4, ORF30-2xStrep pCDNA4, ORF31- 2xStrep pCDNA4, ORF34-2xStrep pCDNA4, and ORF66-2xStrep pCDNA4 have been previously described (8). Plasmid pRL-TK (Promega) was kindly provided by Dr. Russel Vance. Lentiviral packaging plasmids psPAX2 (Addgene plasmid # 12260) and pMD2.G (Addgene plasmid # 12259) were gifts from Dr. Didier Trono.

**Table 1.**
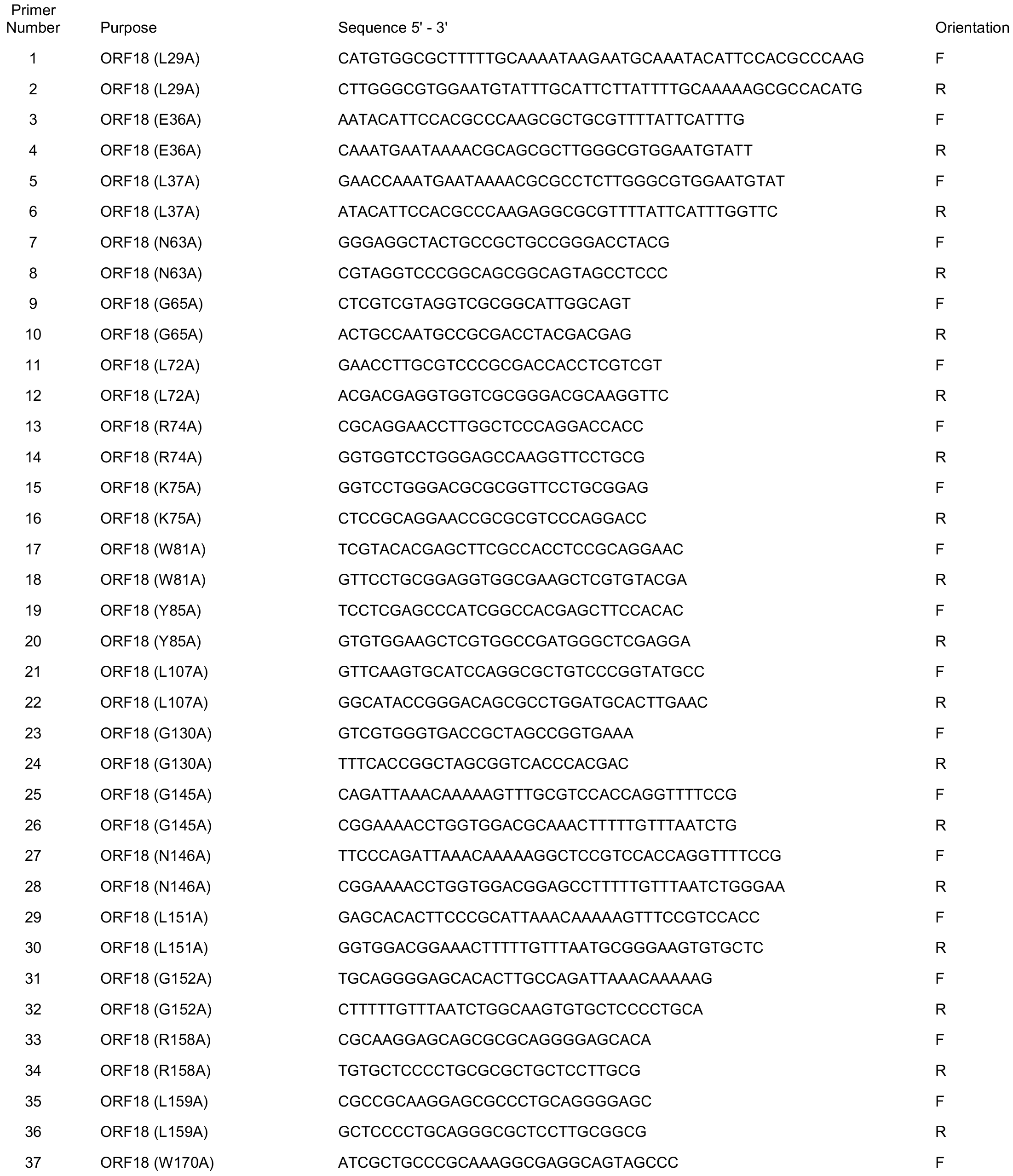
Primer sequences used in this study

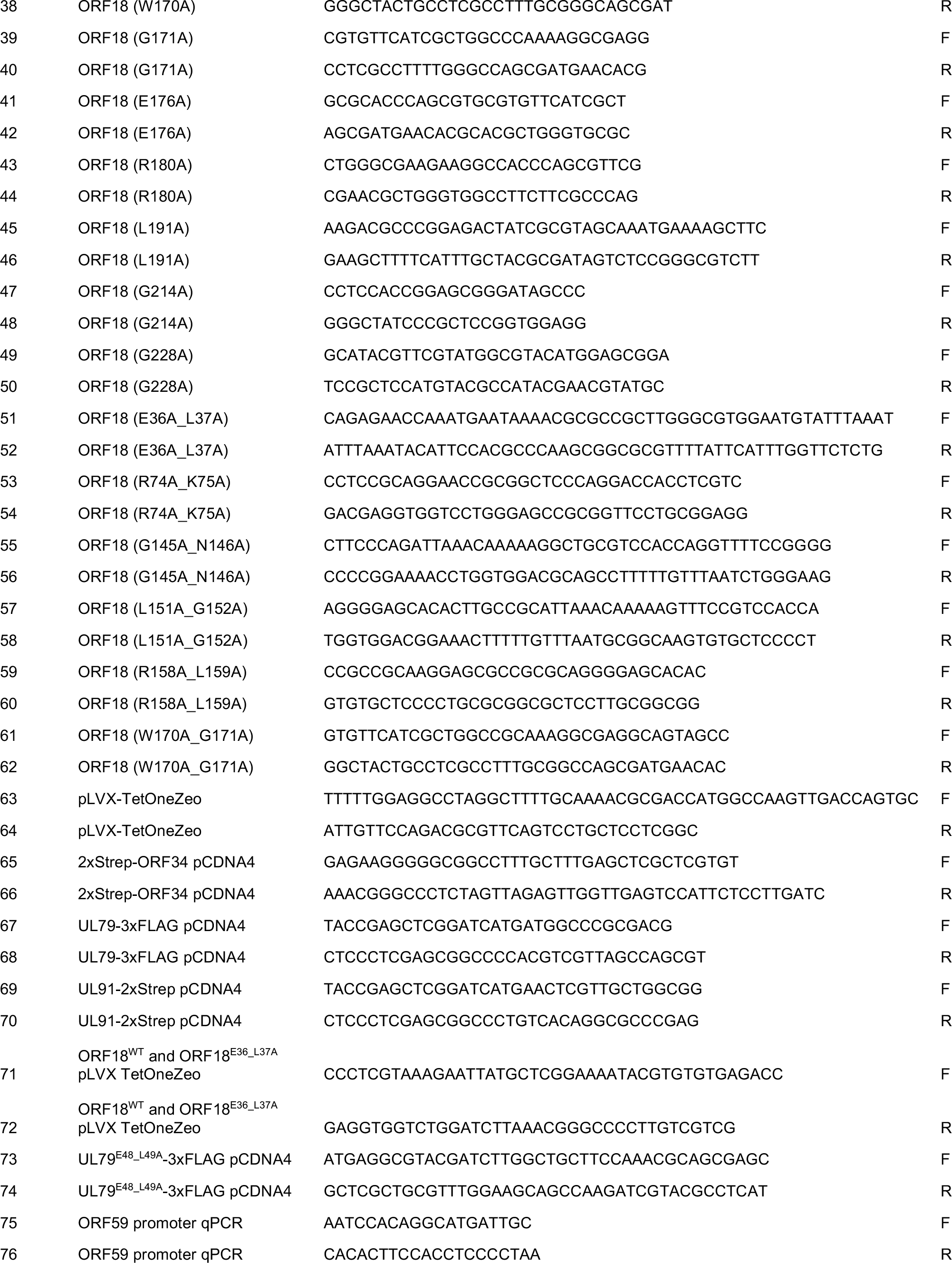

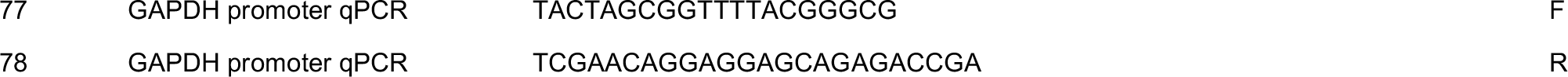

### Cells and transfections

HEK293T cells (ATCC CRL-3216) were maintained in DMEM supplemented with 10% FBS (Seradygm). The iSLK renal carcinoma cell line harboring the KSHV genome on the bacterial artificial chromosome BAC16 and a doxycycline-inducible copy of the KSHV lytic transactivator RTA (iSLK-BAC16) has been described previously (23). iSLK-BAC16-ORF18.stop cells that contain a stop mutation in ORF18 were kindly provided by Dr. Ting-Ting Wu (6). iSLK-BAC16 and iSLK-BAC16-ORF18.stop were maintained in DMEM supplemented with 10% FBS, 1 mg/ml hygromycin, and 1 μg/ml puromycin (iSLK-BAC16 media). iSLK-BAC16-ORF18.stop cells were complemented by lentiviral transduction with ORF18^WT^-3xFLAG or ORF18^E36A_L37A^-3xFLAG. To generate the lentivirus, HEK293T cells were co-transfected with pLVX-TetOneZeo-ORF18^WT^-3xFLAG or pLVX-TetOneZeo-ORF18^E36A_L37A^-3xFLAG along with the packaging plasmids pMD2.G and psPAX2. After 48h, the supernatant was harvested and syringe-filtered through a 0.45 μm filter (Millipore). The supernatant was diluted 1:2 with DMEM and polybrene was added to a final concentration of 8 μg/ml. 1 × 10^6^ iSLK-BAC16-ORF18.stop freshly trypsinized cells were spinfected in a 6-well plate for 2 h at 500 × *g*. After 24 h the cells were expanded to a 10 cm tissue culture plate and selected for 2 weeks in iSLK-BAC16 media supplemented with 325 μg/ml zeocin (Sigma). For DNA transfections, HEK293T cells were plated and transfected after 24 h at 70% confluency with PolyJet (SignaGen).

### Immunoprecipitation and western blotting

Cell lysates were prepared 24 h after transfection by washing and pelleting cells in cold PBS, then resuspending the pellets in IP lysis buffer [50 mM Tris-HCl pH 7.4, 150 mM NaCl, 1mM EDTA, 0.5% NP-40, and protease inhibitor (Roche)] and rotating for 30 minutes at 4 °C. Lysates were cleared by centrifugation at 21,000 × *g* for 10 min, then 1 mg (for pairwise interaction IPs) or 2 mg (for the entire late gene complex IPs) of lysate was incubated with pre-washed M2 anti-FLAG magnetic beads (Sigma) overnight. The beads were washed 3x for 5 min each with IP wash buffer [50 mM Tris-HCl pH 7.4, 150 mM NaCl, 1mM EDTA, 0.05% NP-40] and eluted with 2× Laemmli sample buffer (BioRad).

Lysates and elutions were resolved by SDS-PAGE and western blotted in TBST (Tris-buffered saline, 0.2% Tween 20) using the following primary antibodies: Strep-HRP (Millipore, 1:2500); rabbit anti-FLAG (Sigma, 1:3000); mouse anti-FLAG (Sigma, 1:1000); rabbit anti-Vinculin (Abcam, 1:1000); mouse anti-GAPDH (Abcam, 1:1000); mouse anti-Pol II CTD clone 8WG16 (Abcam, 1:1000); rabbit anti-K8.1 (1:10000); rabbit anti-ORF59 (1:10000). Rabbit anti-ORF59 and anti-K8.1 was produced by the Pocono Rabbit Farm and Laboratory by immunizing rabbits against MBP-ORF59 or MBP-K8.1 [gifts from Denise Whitby (29)]. Following incubation with primary antibodies, the membranes were washed with TBST and incubated with the appropriate secondary antibody. The secondary antibodies used were the following: goat anti-mouse-HRP (1:5000, Southern Biotech) and goat anti-rabbit-HRP (1:5000, Southern Biotech).

The co-IP efficiency for the pairwise interactions was quantified from the western blot images using Image Lab software (BioRad). The band intensity for the both the Strep-tagged ORF and ORF18^WT^-3xFLAG or ORF18^Mu^-3xFLAG was calculated for the IP lanes of the western blot. The ratio of the band intensity of Strep-tagged ORF to ORF18^Mu^-3xFLAG was divided by the ratio of Strep-tagged ORF to ORF18^WT^-3xFLAG to generate a co-IP efficiency for each ORF18^Mu^ relative to the co-IP efficiency of ORF18^WT^.

### ORF30 Protein Stability

Translation was inhibited 24 h after transfection by the addition of 100 μg/ml cycloheximide for 0-8h. Cells were washed once in cold PBS, and cell pellets were frozen until all samples were collected. The pellets were lysed in IP lysis buffer by rotating for 30 minutes at 4 °C. Lysates were cleared by centrifugation at 21,000 × *g* for 10 min, then resolved by SDS-PAGE followed by western blot.

### Virus Characterization

For reactivation studies, 1 × 10^6^ iSLK cells were plated in 10 cm dishes for 16 h, then induced with 1 μg/ml doxycycline and 1 mM sodium butyrate for an additional 72 h. To determine the fold DNA induction in reactivated cells, the cells were scraped and triturated in the induced media, and 200 μl of the cell/supernatant suspension was treated overnight with 80 μg/ml proteinase K (Promega) in 1× proteinase K digestion buffer (10 mM Tris-HCl pH 7.4, 100 mM NaCl, 1 mM EDTA, 0.5% SDS) after which DNA was extracted using a Quick-DNA Miniprep kit (Zymo). Viral DNA fold induction was quantified by qPCR using iTaq Universal SYBR Green Supermix (BioRad) on a QuantStudio3 Real-Time PCR machine with primers 75/76 (Table 1) for the KSHV ORF59 promoter and normalized to the level of GAPDH promoter with primers 77/78 (Table 1).

Infectious virion production was determined by supernatant transfer assay. Supernatant from induced iSLK cells was syringe-filtered through a 0.45 μm filter and diluted 1:2 with DMEM, then 2 mL of the supernatant was spinoculated onto 1 × 10^6^ freshly trypsinized HEK293T cells for 2 h at 500 × *g*. After 24 h, the media was aspirated, the cells were washed once with cold PBS and crosslinked in 4% PFA (Ted Pella) diluted in PBS. The cells were pelleted, resuspended in PBS, and a minimum of 50,000 cells/sample were analyzed on a BD Accuri 6 flow cytometer. The data were analyzed using FlowJo (30).

### Late Gene Reporter Assay

HEK293T cells were plated in 6-well plate and each well was transfected with 900 ng of DNA containing 125 ng each of pcDNA4 ORF18-3xFLAG or ORF18^Mu^-3xFLAG, ORF24-2xStrep, ORF30-2xStrep, ORF31-2xStrep, 2xStrep-ORF34, ORF66-2xStrep, and either K8.1 Pr pGL4.16 or ORF57 Pr pGL4.16, along with 25 ng of pRL-TK as an internal transfection control. After 24 h, the cells were rinsed once with PBS, lysed by rocking for 15 min at room temperature in 500 μl of Passive Lysis Buffer (Promega), and clarified by centrifuging at 21,000 × *g* for 2 min. 20 μl of the clarified lysate was added in triplicate to a white chimney well microplate (Greiner bio-one) to measure luminescence on a Tecan M1000 microplate reader using a Dual Luciferase Assay Kit (Promega). The firefly luminescence was normalized to the internal Renilla luciferase control for each transfection.

## Acknowledgements

We thank all members of the Glaunsinger and Coscoy labs, in particular Matthew R Gardner, for their helpful suggestions and critical reading of the manuscript. This material is based upon work supported by the National Science Foundation Graduate Research Fellowship under Grant No. DGE 1752814 and UC Berkeley Chancellor’s Fellowship awarded to A.C. B.G. is an investigator of the Howard Hughes Medical Institute. This research was also supported by NIH R01AI122528 to B.G.

